## Supplementary material for "High-Performance Classification of Breast Cancer Histopathological Images Using Fine-Tuned Vision Transformers on the BreakHis Dataset": Code: BreaKHis_classifier_final_model_training.ipynb

```
# %% [code]
!pip install transformers datasets evaluate accelerate pillow torchvision scikit-learn

# %% [code]
from huggingface_hub import login
login("your login api token")

from google.colab import drive
drive.mount('/content/drive')

# %% [code]
import torch

torch.cuda.empty_cache()

# %% [code]
!tar -xvf /content/drive/MyDrive/breast_histology/BreaKHis_v1.tar.gz

# %% [code]
###written on command by CHATGPT4

import os
import shutil

# Define the base directory for the current dataset structure
base_dir = 'BreaKHis_v1/histology_slides/breast'

# Define the target base directory for the new structure
target_base_dir = 'reorganized_breakhis'

# Define the zoom levels you are interested in
zoom_levels = ['40X', '100X', '200X', '400X']

# Create the target base directory if it doesn't exist
os.makedirs(target_base_dir, exist_ok=True)

benign_dir = os.path.join(target_base_dir, 'benign')
malignant_dir = os.path.join(target_base_dir, 'malignant')
os.makedirs(benign_dir)
os.makedirs(malignant_dir)

for zoom in zoom_levels:
    # Create directories for each zoom level under the target base directory

    # Create subdirectories for benign and malignant under each zoom level directory


    # Iterate through the malignant and benign directories in the current dataset
    for category in ['malignant', 'benign']:
        category_dir = os.path.join(base_dir, category)

        # Walk through the directory structure
        for root, dirs, files in os.walk(category_dir):
            # Check if the current directory is the desired zoom level directory
            if os.path.basename(root) == zoom:
                # Determine the target directory based on the category (benign or malignant)
                target_dir = malignant_dir if category == 'malignant' else benign_dir

                # Move all image files to the target directory
                for file in files:
                    if file.endswith('.png'):
                        source_file = os.path.join(root, file)
                        target_file = os.path.join(target_dir, file)
                        shutil.move(source_file, target_file)

print("Dataset reorganization complete!")

# %% [code]
from datasets import load_dataset
breakhis = load_dataset("imagefolder", data_dir=target_base_dir, split='train')


# %% [code]
#run this for 5 fold crossval
from sklearn.model_selection import KFold

kf = KFold(n_splits=5, shuffle=True)
splits = []

# Generate indices for the splits
for train_index, test_index in kf.split(breakhis):
    # Split the dataset based on the indices
    train_split = breakhis.select(train_index)
    test_split = breakhis.select(test_index)
    splits.append((train_split, test_split))


# %% [code]
print(len(splits))
print((splits[0][1]))

# %% [code]
labels=['benign', 'malignant']
label2id, id2label= dict(), dict()

for i, label in enumerate(labels):
  label2id[label] = str(i)
  id2label[str(i)] = label

# %% [code]
from transformers import AutoImageProcessor
from torchvision.transforms import RandomResizedCrop, Compose, Normalize, ToTensor

checkpoint="google/vit-base-patch16-224-in21k"

image_processor=AutoImageProcessor.from_pretrained(checkpoint)


normalize = Normalize(mean=image_processor.image_mean, std=image_processor.image_std)

size = (
    image_processor.size["shortest_edge"]
    if "shortest_edge" in image_processor.size
    else (image_processor.size["height"], image_processor.size["width"])
)

_transforms = Compose([RandomResizedCrop(size), ToTensor(), normalize])

def transforms(examples):
    examples["pixel_values"] = [_transforms(img.convert("RGB")) for img in examples["image"]]
    del examples["image"]
    return examples

# %% [code]
from tensorflow import keras
from tensorflow.keras import layers

size = (image_processor.size["height"], image_processor.size["width"])

train_data_augmentation = keras.Sequential(
    [
        layers.RandomCrop(size[0], size[1]),
        layers.Rescaling(scale=1.0 / 127.5, offset=-1),
        layers.RandomFlip("horizontal"),
        layers.RandomRotation(factor=0.02),
        layers.RandomZoom(height_factor=0.2, width_factor=0.2),
    ],
    name="train_data_augmentation",
)

val_data_augmentation = keras.Sequential(
    [
        layers.CenterCrop(size[0], size[1]),
        layers.Rescaling(scale=1.0 / 127.5, offset=-1),
    ],
    name="val_data_augmentation",
)

# %% [code]
import numpy as np
import tensorflow as tf
from PIL import Image


def convert_to_tf_tensor(image: Image):
    np_image = np.array(image)
    tf_image = tf.convert_to_tensor(np_image)
    # `expand_dims()` is used to add a batch dimension since
    # the TF augmentation layers operates on batched inputs.
    return tf.expand_dims(tf_image, 0)


def preprocess_train(example_batch):
    """Apply train_transforms across a batch."""
    images = [
        train_data_augmentation(convert_to_tf_tensor(image.convert("RGB"))) for image in example_batch["image"]
    ]
    example_batch["pixel_values"] = [tf.transpose(tf.squeeze(image)) for image in images]
    return example_batch


def preprocess_val(example_batch):
    """Apply val_transforms across a batch."""
    images = [
        val_data_augmentation(convert_to_tf_tensor(image.convert("RGB"))) for image in example_batch["image"]
    ]
    example_batch["pixel_values"] = [tf.transpose(tf.squeeze(image)) for image in images]
    return example_batch

# %% [code]
print(splits[0][0])

# %% [code]
for i in splits:
  i[0].set_transform(preprocess_train)
  i[1].set_transform(preprocess_val)

# %% [code]
splits_transforms = []
for i in splits:
  cur = []
  cur.append(i[0].with_transform(transforms))
  cur.append(i[1].with_transform(transforms))
  splits_transforms.append(cur)


# %% [code]
len(splits_transforms)

# %% [code]
from transformers import DefaultDataCollator

data_collator = DefaultDataCollator()

# %% [code]
from transformers import AutoModelForImageClassification, TrainingArguments, Trainer

model = AutoModelForImageClassification.from_pretrained(
    checkpoint,
    num_labels=len(labels),
    id2label=id2label,
    label2id=label2id,
)

# %% [code]
#ranger class as implemented in Wright et al. 2021, "Ranger21: A synergistic deep learning optimizer"
import math
import torch
from torch.optim.optimizer import Optimizer

torch.cuda.empty_cache()

class Ranger(Optimizer):

	def __init__(self, params, lr=1e-3,  # lr
				 alpha=0.5, k=6, N_sma_threshhold=5,  # Ranger options
				 betas=(.95, 0.999), eps=1e-5, weight_decay=0,  # Adam options
				 use_gc=True, gc_conv_only=False
				 # Gradient centralization on or off, applied to conv layers only or conv + fc layers
				 ):

		# parameter checks
		if not 0.0 <= alpha <= 1.0:
			raise ValueError(f'Invalid slow update rate: {alpha}')
		if not 1 <= k:
			raise ValueError(f'Invalid lookahead steps: {k}')
		if not lr > 0:
			raise ValueError(f'Invalid Learning Rate: {lr}')
		if not eps > 0:
			raise ValueError(f'Invalid eps: {eps}')

		# parameter comments:
		# beta1 (momentum) of .95 seems to work better than .90...
		# N_sma_threshold of 5 seems better in testing than 4.
		# In both cases, worth testing on your dataset (.90 vs .95, 4 vs 5) to make sure which works best for you.

		# prep defaults and init torch.optim base
		defaults = dict(lr=lr, alpha=alpha, k=k, step_counter=0, betas=betas, N_sma_threshhold=N_sma_threshhold,
						eps=eps, weight_decay=weight_decay)
		super().__init__(params, defaults)

		# adjustable threshold
		self.N_sma_threshhold = N_sma_threshhold

		# look ahead params

		self.alpha = alpha
		self.k = k

		# radam buffer for state
		self.radam_buffer = [[None, None, None] for ind in range(10)]

		# gc on or off
		self.use_gc = use_gc

		# level of gradient centralization
		self.gc_gradient_threshold = 3 if gc_conv_only else 1

	def __setstate__(self, state):
		super(Ranger, self).__setstate__(state)

	def step(self, closure=None):
		loss = None

		# Evaluate averages and grad, update param tensors
		for group in self.param_groups:

			for p in group['params']:
				if p.grad is None:
					continue
				grad = p.grad.data.float()

				if grad.is_sparse:
					raise RuntimeError('Ranger optimizer does not support sparse gradients')

				p_data_fp32 = p.data.float()

				state = self.state[p]  # get state dict for this param

				if len(state) == 0:  # if first time to run...init dictionary with our desired entries
					# if self.first_run_check==0:
					# self.first_run_check=1
					# print("Initializing slow buffer...should not see this at load from saved model!")
					state['step'] = 0
					state['exp_avg'] = torch.zeros_like(p_data_fp32)
					state['exp_avg_sq'] = torch.zeros_like(p_data_fp32)

					# look ahead weight storage now in state dict
					state['slow_buffer'] = torch.empty_like(p.data)
					state['slow_buffer'].copy_(p.data)

				else:
					state['exp_avg'] = state['exp_avg'].type_as(p_data_fp32)
					state['exp_avg_sq'] = state['exp_avg_sq'].type_as(p_data_fp32)

				# begin computations
				exp_avg, exp_avg_sq = state['exp_avg'], state['exp_avg_sq']
				beta1, beta2 = group['betas']

				# GC operation for Conv layers and FC layers
				if grad.dim() > self.gc_gradient_threshold:
					grad.add_(-grad.mean(dim=tuple(range(1, grad.dim())), keepdim=True))

				state['step'] += 1

				# compute variance mov avg
				exp_avg_sq.mul_(beta2).addcmul_(1 - beta2, grad, grad)
				# compute mean moving avg
				exp_avg.mul_(beta1).add_(1 - beta1, grad)

				buffered = self.radam_buffer[int(state['step'] % 10)]

				if state['step'] == buffered[0]:
					N_sma, step_size = buffered[1], buffered[2]
				else:
					buffered[0] = state['step']
					beta2_t = beta2 ** state['step']
					N_sma_max = 2 / (1 - beta2) - 1
					N_sma = N_sma_max - 2 * state['step'] * beta2_t / (1 - beta2_t)
					buffered[1] = N_sma
					if N_sma > self.N_sma_threshhold:
						step_size = math.sqrt(
							(1 - beta2_t) * (N_sma - 4) / (N_sma_max - 4) * (N_sma - 2) / N_sma * N_sma_max / (
										N_sma_max - 2)) / (1 - beta1 ** state['step'])
					else:
						step_size = 1.0 / (1 - beta1 ** state['step'])
					buffered[2] = step_size

				if group['weight_decay'] != 0:
					p_data_fp32.add_(-group['weight_decay'] * group['lr'], p_data_fp32)

				# apply lr
				if N_sma > self.N_sma_threshhold:
					denom = exp_avg_sq.sqrt().add_(group['eps'])
					p_data_fp32.addcdiv_(-step_size * group['lr'], exp_avg, denom)
				else:
					p_data_fp32.add_(-step_size * group['lr'], exp_avg)

				p.data.copy_(p_data_fp32)

				# integrated look ahead...
				# we do it at the param level instead of group level
				if state['step'] % group['k'] == 0:
					slow_p = state['slow_buffer']  # get access to slow param tensor
					slow_p.add_(self.alpha, p.data - slow_p)  # (fast weights - slow weights) * alpha
					p.data.copy_(slow_p)  # copy interpolated weights to RAdam param tensor

		return loss

# %% [code]
import evaluate

accuracy = evaluate.load("accuracy")

import numpy as np

def compute_metrics(eval_pred):
    predictions, labels = eval_pred
    predictions = np.argmax(predictions, axis=1)
    accuracy_metric = accuracy.compute(predictions=predictions, references=labels)
    return accuracy_metric

# %% [code]
from transformers import get_scheduler
import pandas as pd

optimizer = Ranger(model.parameters(), lr=3e-5, weight_decay=.02)
training_args = TrainingArguments(
    output_dir="break_his_default_ranger_rotations_5_fold_1_final",
    remove_unused_columns=False,
    eval_strategy="epoch",
    save_strategy="epoch",
    learning_rate=3e-5,
    per_device_train_batch_size=32,
    gradient_accumulation_steps=8,
    per_device_eval_batch_size=32,
    num_train_epochs=100,
    warmup_ratio=0.2,
    logging_steps=10,
    load_best_model_at_end=False,
    metric_for_best_model="accuracy",
    push_to_hub=True,
    hub_private_repo=True,
    hub_model_id="sloshywings/breakhis-model-ranger-rotations-5-fold-1-final"
)

training_steps = training_args.num_train_epochs * len(splits_transforms[0][0]) // training_args.per_device_train_batch_size
lr_scheduler = get_scheduler(
    name="cosine",
    optimizer=optimizer,
    num_warmup_steps=training_steps//6
    ,
    num_training_steps=training_steps
)

for i in splits_transforms:
    test_dataset = i[1]
  # Create a Trainer instance
    train_dataset = i[0]
    trainer = Trainer(
        model=model,
        args=training_args,
        data_collator=data_collator,
        train_dataset=train_dataset,
        eval_dataset=test_dataset,
        optimizers=(optimizer, lr_scheduler),
        tokenizer=image_processor,
        compute_metrics=compute_metrics,
    )

    trainer.train()

res_log = pd.DataFrame(trainer.state.log_history)


# %% [code]
trainer.push_to_hub()
```
