## Supplementary material for "High-Performance Classification of Breast Cancer Histopathological Images Using Fine-Tuned Vision Transformers on the BreakHis Dataset": Code: BreaKHis_final_classifier_models_eval.ipynb

```
# %% [code]
!pip install transformers datasets evaluate accelerate pillow torchvision scikit-learn

# %% [code]
from huggingface_hub import login
login("enter your token with read and write permissions")

from google.colab import drive
drive.mount('/content/drive')

# %% [code]
import torch

torch.cuda.empty_cache()

# %% [code]
!tar -xvf /content/drive/MyDrive/breast_histology/BreaKHis_v1.tar.gz

# %% [code]
###written in part on command by CHATGPT4
#prepare breakhis dataset for accuracy check
import os
import shutil

# Define the base directory for the current dataset structure
base_dir = 'BreaKHis_v1/histology_slides/breast'

# Define the target base directory for the new structure
target_base_dir = 'reorganized_breakhis'

# Define the zoom levels you are interested in
zoom_levels = ['40X', '100X', '200X', '400X']

# Create the target base directory if it doesn't exist
os.makedirs(target_base_dir, exist_ok=True)

benign_dir = os.path.join(target_base_dir, 'benign')
malignant_dir = os.path.join(target_base_dir, 'malignant')
os.makedirs(benign_dir)
os.makedirs(malignant_dir)

for zoom in zoom_levels:
    # Create directories for each zoom level under the target base directory

    # Create subdirectories for benign and malignant under each zoom level directory


    # Iterate through the malignant and benign directories in the current dataset
    for category in ['malignant', 'benign']:
        category_dir = os.path.join(base_dir, category)

        # Walk through the directory structure
        for root, dirs, files in os.walk(category_dir):
            # Check if the current directory is the desired zoom level directory
            if os.path.basename(root) == zoom:
                # Determine the target directory based on the category (benign or malignant)
                target_dir = malignant_dir if category == 'malignant' else benign_dir

                # Move all image files to the target directory
                for file in files:
                    if file.endswith('.png'):
                        source_file = os.path.join(root, file)
                        target_file = os.path.join(target_dir, file)
                        shutil.move(source_file, target_file)

print("Dataset reorganization complete!")

# %% [code]
#load base model for either transfer learning or accuracy check
import torch
from transformers import AutoModelForImageClassification, AutoFeatureExtractor
from torchvision.datasets import ImageFolder
from torchvision.transforms import Compose, Normalize, Resize, ToTensor
from torch.utils.data import DataLoader
from sklearn.metrics import confusion_matrix, precision_score, recall_score, f1_score, accuracy_score, roc_curve, matthews_corrcoef

# Load the pre-trained model and feature extractor
#model_id = "sloshywings/breakhis-model-ranger-alt-retrial-1"
#model_id = "sloshywings/breakhis-model-ranger-rotations-0"
model_id="sloshywings/breakhis-model-ranger-rotations-5-fold-1-final"
print(model_id)
model = AutoModelForImageClassification.from_pretrained(model_id)
feature_extractor = AutoFeatureExtractor.from_pretrained(model_id)

# Define transformations
transform = Compose([
    Resize((224, 224)),
    ToTensor(),
    Normalize(mean=feature_extractor.image_mean, std=feature_extractor.image_std)
])


# %% [code]
#check accuracy and statistics for breakhis dataset
# Load the dataset using ImageFolder
dataset = ImageFolder(root='reorganized_breakhis', transform=transform)

# Create DataLoader
data_loader = DataLoader(dataset, batch_size=328, shuffle=False)

# Move model to GPU if available
device = torch.device('cuda' if torch.cuda.is_available() else 'cpu')
model.to(device)
model.eval()

# Initialize lists to store targets and predictions
true_labels = []
pred_labels = []

# Predict the labels for the test set
with torch.no_grad():
    for batch in data_loader:
        inputs, labels = batch
        inputs = inputs.to(device)
        labels = labels.to(device)
        outputs = model(inputs)
        logits = outputs.logits
        predictions = torch.argmax(logits, dim=-1)
        true_labels.extend(labels.cpu().numpy())
        pred_labels.extend(predictions.cpu().numpy())

# Calculate metrics
def calculate_metrics(y_true, y_pred):
    accuracy = accuracy_score(y_true, y_pred)
    roc = roc_curve(y_true, y_pred)
    precision = precision_score(y_true, y_pred, average='macro')
    recall = recall_score(y_true, y_pred, average='macro')
    f1 = f1_score(y_true, y_pred, average='macro')
    tn, fp, fn, tp = confusion_matrix(y_true, y_pred).ravel()
    specificity = tn / (tn + fp)
    fdr = fp / (tp + fp)
    fnr = fn / (fn + tp)
    fpr = fp / (fp + tn)
    mcc = matthews_corrcoef(y_true, y_pred)
    npv = tn / (tn + fn)

    return {
        "Accuracy": accuracy,
        "Precision": precision,
        "Recall": recall,
        "F1 Score": f1,
        "Specificity": specificity,
        "FDR": fdr,
        "FNR": fnr,
        "FPR": fpr,
        "MCC": mcc,
        "NPV": npv,
        "roc": roc
    }

# Calculate and print all metrics
metrics = calculate_metrics(true_labels, pred_labels)

for metric, value in metrics.items():
    try:
      print(f"{metric}: {value:.4f}")
    except:
      print(f"{metric}: {value}")


# %% [code]
!unzip /content/drive/MyDrive/breast_histology/BACH.zip

# %% [code]
#generate dataset for BACH
import shutil
import os

# Define the source and destination directories
src_b = "/content/ICIAR2018_BACH_Challenge/ICIAR2018_BACH_Challenge/Photos/Benign"
src_m1 = "/content/ICIAR2018_BACH_Challenge/ICIAR2018_BACH_Challenge/Photos/Invasive"
#src_m2 = "/content/ICIAR2018_BACH_Challenge/ICIAR2018_BACH_Challenge/Photos/InSitu"

#IN SITU IS NOT MALIGNANT
all_src_m = [src_m1]

dest_m = 'reorganized_BACH/malignant'
dest_b = 'reorganized_BACH/benign'

# Create the destination directory if it doesn't exist
os.makedirs('reorganized_BACH', exist_ok=True)

os.makedirs(dest_m, exist_ok=True)

os.makedirs(dest_b, exist_ok=True)
# Copy all files from the source directory to the destination directory
for file_name in os.listdir(src_b):
    source_file = os.path.join(src_b, file_name)
    destination_file = os.path.join(dest_b, file_name)
    if os.path.isfile(source_file):
        shutil.copy(source_file, destination_file)

print(f"All files have been copied from '{src_b}' to '{dest_b}'.")

for cur_src_m in all_src_m:
  for file_name in os.listdir(cur_src_m):
    source_file = os.path.join(cur_src_m, file_name)
    destination_file = os.path.join(dest_m, file_name)
    if os.path.isfile(source_file):
        shutil.copy(source_file, destination_file)

  print(f"All files have been copied from '{cur_src_m}' to '{dest_m}'.")


# %% [code]
#accuracy check for transfer learning model on BACH dataset
import torch
from transformers import AutoModelForImageClassification, AutoFeatureExtractor
from torchvision.datasets import ImageFolder
from torchvision.transforms import Compose, Normalize, Resize, ToTensor
from torch.utils.data import DataLoader
from sklearn.metrics import confusion_matrix, precision_score, recall_score, f1_score, accuracy_score, roc_curve, matthews_corrcoef

model_id="sloshywings/breakhis-model-ranger-rotations-5-fold-1-transfer-learning-bach-final"

print(model_id)
model = AutoModelForImageClassification.from_pretrained(model_id)
feature_extractor = AutoFeatureExtractor.from_pretrained(model_id)

# Define transformations
transform = Compose([
    Resize((224, 224)),
    ToTensor(),
    Normalize(mean=feature_extractor.image_mean, std=feature_extractor.image_std)
])

# Load the dataset using ImageFolder
dataset = ImageFolder(root='reorganized_BACH', transform=transform)
#dataset = ImageFolder(root='reorganized_idc', transform=transform)

# Create DataLoader
data_loader = DataLoader(dataset, batch_size=328, shuffle=False)

# Move model to GPU if available
device = torch.device('cuda' if torch.cuda.is_available() else 'cpu')
model.to(device)
model.eval()

# Initialize lists to store targets and predictions
true_labels = []
pred_labels = []

# Predict the labels for the test set
with torch.no_grad():
    for batch in data_loader:
        inputs, labels = batch
        inputs = inputs.to(device)
        labels = labels.to(device)
        outputs = model(inputs)
        logits = outputs.logits
        predictions = torch.argmax(logits, dim=-1)
        true_labels.extend(labels.cpu().numpy())
        pred_labels.extend(predictions.cpu().numpy())

# Calculate metrics
def calculate_metrics(y_true, y_pred):
    accuracy = accuracy_score(y_true, y_pred)
    roc = roc_curve(y_true, y_pred)
    precision = precision_score(y_true, y_pred, average='macro')
    recall = recall_score(y_true, y_pred, average='macro')
    f1 = f1_score(y_true, y_pred, average='macro')
    tn, fp, fn, tp = confusion_matrix(y_true, y_pred).ravel()
    specificity = tn / (tn + fp)
    fdr = fp / (tp + fp)
    fnr = fn / (fn + tp)
    fpr = fp / (fp + tn)
    mcc = matthews_corrcoef(y_true, y_pred)
    npv = tn / (tn + fn)

    return {
        "Accuracy": accuracy,
        "Precision": precision,
        "Recall": recall,
        "F1 Score": f1,
        "Specificity": specificity,
        "FDR": fdr,
        "FNR": fnr,
        "FPR": fpr,
        "MCC": mcc,
        "NPV": npv,
        "roc": roc
    }

# Calculate and print all metrics
metrics = calculate_metrics(true_labels, pred_labels)

for metric, value in metrics.items():
    try:
      print(f"{metric}: {value:.4f}")
    except:
      print(f"{metric}: {value}")


# %% [code]
!mkdir /content/IDC_dataset/
!unzip /content/drive/MyDrive/breast_histology/IDC_dataset.zip -d /content/IDC_dataset/

# %% [code]
#Generate dataset for IDC
import os
import shutil

# Path to the root directory containing the numbered folders
root_dir = 'IDC_dataset/'

# Paths for the new directories to be created
benign_dir = os.path.join('reorganized_idc/benign')
malignant_dir = os.path.join('reorganized_idc/malignant')

os.makedirs('reogranized_idc/', exist_ok=True)

# Create the new directories if they don't exist
os.makedirs(benign_dir, exist_ok=True)
os.makedirs(malignant_dir, exist_ok=True)

# Iterate over each numbered folder in the root directory
for folder in os.listdir(root_dir):
    folder_path = os.path.join(root_dir, folder)

    # Check if it's a directory
    if os.path.isdir(folder_path):
        # Define the paths for the 0 (benign) and 1 (malignant) folders
        benign_folder_path = os.path.join(folder_path, '0')
        malignant_folder_path = os.path.join(folder_path, '1')

        # If the benign folder exists, copy its contents to the new benign directory
        if os.path.exists(benign_folder_path):
            for file_name in os.listdir(benign_folder_path):
                shutil.copy(os.path.join(benign_folder_path, file_name), benign_dir)

        # If the malignant folder exists, copy its contents to the new malignant directory
        if os.path.exists(malignant_folder_path):
            for file_name in os.listdir(malignant_folder_path):
                shutil.copy(os.path.join(malignant_folder_path, file_name), malignant_dir)


# %% [code]
#ranger class as implemented in Wright et al. 2021, "Ranger21: A synergistic deep learning optimizer"
import math
import torch
from torch.optim.optimizer import Optimizer

torch.cuda.empty_cache()

class Ranger(Optimizer):

	def __init__(self, params, lr=1e-3,  # lr
				 alpha=0.5, k=6, N_sma_threshhold=5,  # Ranger options
				 betas=(.95, 0.999), eps=1e-5, weight_decay=0,  # Adam options
				 use_gc=True, gc_conv_only=False
				 # Gradient centralization on or off, applied to conv layers only or conv + fc layers
				 ):

		# parameter checks
		if not 0.0 <= alpha <= 1.0:
			raise ValueError(f'Invalid slow update rate: {alpha}')
		if not 1 <= k:
			raise ValueError(f'Invalid lookahead steps: {k}')
		if not lr > 0:
			raise ValueError(f'Invalid Learning Rate: {lr}')
		if not eps > 0:
			raise ValueError(f'Invalid eps: {eps}')

		# parameter comments:
		# beta1 (momentum) of .95 seems to work better than .90...
		# N_sma_threshold of 5 seems better in testing than 4.
		# In both cases, worth testing on your dataset (.90 vs .95, 4 vs 5) to make sure which works best for you.

		# prep defaults and init torch.optim base
		defaults = dict(lr=lr, alpha=alpha, k=k, step_counter=0, betas=betas, N_sma_threshhold=N_sma_threshhold,
						eps=eps, weight_decay=weight_decay)
		super().__init__(params, defaults)

		# adjustable threshold
		self.N_sma_threshhold = N_sma_threshhold

		# look ahead params

		self.alpha = alpha
		self.k = k

		# radam buffer for state
		self.radam_buffer = [[None, None, None] for ind in range(10)]

		# gc on or off
		self.use_gc = use_gc

		# level of gradient centralization
		self.gc_gradient_threshold = 3 if gc_conv_only else 1

	def __setstate__(self, state):
		super(Ranger, self).__setstate__(state)

	def step(self, closure=None):
		loss = None

		# Evaluate averages and grad, update param tensors
		for group in self.param_groups:

			for p in group['params']:
				if p.grad is None:
					continue
				grad = p.grad.data.float()

				if grad.is_sparse:
					raise RuntimeError('Ranger optimizer does not support sparse gradients')

				p_data_fp32 = p.data.float()

				state = self.state[p]  # get state dict for this param

				if len(state) == 0:  # if first time to run...init dictionary with our desired entries
					# if self.first_run_check==0:
					# self.first_run_check=1
					# print("Initializing slow buffer...should not see this at load from saved model!")
					state['step'] = 0
					state['exp_avg'] = torch.zeros_like(p_data_fp32)
					state['exp_avg_sq'] = torch.zeros_like(p_data_fp32)

					# look ahead weight storage now in state dict
					state['slow_buffer'] = torch.empty_like(p.data)
					state['slow_buffer'].copy_(p.data)

				else:
					state['exp_avg'] = state['exp_avg'].type_as(p_data_fp32)
					state['exp_avg_sq'] = state['exp_avg_sq'].type_as(p_data_fp32)

				# begin computations
				exp_avg, exp_avg_sq = state['exp_avg'], state['exp_avg_sq']
				beta1, beta2 = group['betas']

				# GC operation for Conv layers and FC layers
				if grad.dim() > self.gc_gradient_threshold:
					grad.add_(-grad.mean(dim=tuple(range(1, grad.dim())), keepdim=True))

				state['step'] += 1

				# compute variance mov avg
				exp_avg_sq.mul_(beta2).addcmul_(1 - beta2, grad, grad)
				# compute mean moving avg
				exp_avg.mul_(beta1).add_(1 - beta1, grad)

				buffered = self.radam_buffer[int(state['step'] % 10)]

				if state['step'] == buffered[0]:
					N_sma, step_size = buffered[1], buffered[2]
				else:
					buffered[0] = state['step']
					beta2_t = beta2 ** state['step']
					N_sma_max = 2 / (1 - beta2) - 1
					N_sma = N_sma_max - 2 * state['step'] * beta2_t / (1 - beta2_t)
					buffered[1] = N_sma
					if N_sma > self.N_sma_threshhold:
						step_size = math.sqrt(
							(1 - beta2_t) * (N_sma - 4) / (N_sma_max - 4) * (N_sma - 2) / N_sma * N_sma_max / (
										N_sma_max - 2)) / (1 - beta1 ** state['step'])
					else:
						step_size = 1.0 / (1 - beta1 ** state['step'])
					buffered[2] = step_size

				if group['weight_decay'] != 0:
					p_data_fp32.add_(-group['weight_decay'] * group['lr'], p_data_fp32)

				# apply lr
				if N_sma > self.N_sma_threshhold:
					denom = exp_avg_sq.sqrt().add_(group['eps'])
					p_data_fp32.addcdiv_(-step_size * group['lr'], exp_avg, denom)
				else:
					p_data_fp32.add_(-step_size * group['lr'], exp_avg)

				p.data.copy_(p_data_fp32)

				# integrated look ahead...
				# we do it at the param level instead of group level
				if state['step'] % group['k'] == 0:
					slow_p = state['slow_buffer']  # get access to slow param tensor
					slow_p.add_(self.alpha, p.data - slow_p)  # (fast weights - slow weights) * alpha
					p.data.copy_(slow_p)  # copy interpolated weights to RAdam param tensor

		return loss

import evaluate

accuracy = evaluate.load("accuracy")

import numpy as np

def compute_metrics(eval_pred):
    predictions, labels = eval_pred
    predictions = np.argmax(predictions, axis=1)
    accuracy_metric = accuracy.compute(predictions=predictions, references=labels)
    return accuracy_metric

# %% [code]
model_id="sloshywings/breakhis-model-ranger-rotations-5-fold-1-transfer-learning-idc-final"

# %% [code]
#accuracy check for transfer learning model on IDC dataset
import torch
from transformers import AutoModelForImageClassification, AutoFeatureExtractor
from torchvision.datasets import ImageFolder
from torchvision.transforms import Compose, Normalize, Resize, ToTensor
from torch.utils.data import DataLoader
from sklearn.metrics import confusion_matrix, precision_score, recall_score, f1_score, accuracy_score, roc_curve, matthews_corrcoef

model_id="sloshywings/breakhis-model-ranger-rotations-5-fold-1-transfer-learning-idc-final"

print(model_id)
model = AutoModelForImageClassification.from_pretrained(model_id)
feature_extractor = AutoFeatureExtractor.from_pretrained(model_id)

# Define transformations
transform = Compose([
    Resize((224, 224)),
    ToTensor(),
    Normalize(mean=feature_extractor.image_mean, std=feature_extractor.image_std)
])

# Load the dataset using ImageFolder
dataset = ImageFolder(root='reorganized_idc', transform=transform)

# Create DataLoader
data_loader = DataLoader(dataset, batch_size=328, shuffle=False)

# Move model to GPU if available
device = torch.device('cuda' if torch.cuda.is_available() else 'cpu')
model.to(device)
model.eval()

# Initialize lists to store targets and predictions
true_labels = []
pred_labels = []

# Predict the labels for the test set
with torch.no_grad():
    for batch in data_loader:
        inputs, labels = batch
        inputs = inputs.to(device)
        labels = labels.to(device)
        outputs = model(inputs)
        logits = outputs.logits
        predictions = torch.argmax(logits, dim=-1)
        true_labels.extend(labels.cpu().numpy())
        pred_labels.extend(predictions.cpu().numpy())

# Calculate metrics
def calculate_metrics(y_true, y_pred):
    accuracy = accuracy_score(y_true, y_pred)
    roc = roc_curve(y_true, y_pred)
    precision = precision_score(y_true, y_pred, average='macro')
    recall = recall_score(y_true, y_pred, average='macro')
    f1 = f1_score(y_true, y_pred, average='macro')
    tn, fp, fn, tp = confusion_matrix(y_true, y_pred).ravel()
    specificity = tn / (tn + fp)
    fdr = fp / (tp + fp)
    fnr = fn / (fn + tp)
    fpr = fp / (fp + tn)
    mcc = matthews_corrcoef(y_true, y_pred)
    npv = tn / (tn + fn)

    return {
        "Accuracy": accuracy,
        "Precision": precision,
        "Recall": recall,
        "F1 Score": f1,
        "Specificity": specificity,
        "FDR": fdr,
        "FNR": fnr,
        "FPR": fpr,
        "MCC": mcc,
        "NPV": npv,
        "roc": roc
    }

# Calculate and print all metrics
metrics = calculate_metrics(true_labels, pred_labels)

for metric, value in metrics.items():
    try:
      print(f"{metric}: {value:.4f}")
    except:
      print(f"{metric}: {value}")
```
