## Supplementary material for "High-Performance Classification of Breast Cancer Histopathological Images Using Fine-Tuned Vision Transformers on the BreakHis Dataset": Code: breakhis_final_classifier_transfer_learning_model_training.py

```
# -*- coding: utf-8 -*-
"""BreaKHis_final_classifier_transfer_learning_model_training

Automatically generated by Colab.

There are import statements throughout the file since this was modified from the notebook files

Original file was run on google colab. The colab notebooks as run are also included in the python_notebooks directory.  
"""

# Library requirements: pip install transformers datasets evaluate accelerate pillow torchvision scikit-learn

from huggingface_hub import login
login("ENTER HUGGINGFACE API TOKEN WITH READ AND WRITE PERMISSIONS")

#from google.colab import drive
#drive.mount('/content/drive')

import torch

torch.cuda.empty_cache()

#load base model for either transfer learning or accuracy check
import torch
from transformers import AutoModelForImageClassification, AutoFeatureExtractor
from torchvision.datasets import ImageFolder
from torchvision.transforms import Compose, Normalize, Resize, ToTensor
from torch.utils.data import DataLoader
from sklearn.metrics import confusion_matrix, precision_score, recall_score, f1_score, accuracy_score, roc_curve, matthews_corrcoef

# Load the pre-trained model and feature extractor

model_id="sloshywings/breakhis-model-ranger-rotations-5-fold-0"

print(model_id)
model = AutoModelForImageClassification.from_pretrained(model_id)
feature_extractor = AutoFeatureExtractor.from_pretrained(model_id)

# Define transformations
transform = Compose([
    Resize((224, 224)),
    ToTensor(),
    Normalize(mean=feature_extractor.image_mean, std=feature_extractor.image_std)
])

#This file assumes that the BACH.zip dataset is unpacked as described in this command at the current level directory: unzip /content/drive/MyDrive/breast_histology/BACH.zip

#generate dataset for BACH
import shutil
import os

# Define the source and destination directories
src_b = "/content/ICIAR2018_BACH_Challenge/ICIAR2018_BACH_Challenge/Photos/Benign"
src_m1 = "/content/ICIAR2018_BACH_Challenge/ICIAR2018_BACH_Challenge/Photos/Invasive"
#src_m2 = "/content/ICIAR2018_BACH_Challenge/ICIAR2018_BACH_Challenge/Photos/InSitu"

#IN SITU IS NOT MALIGNANT
all_src_m = [src_m1]

dest_m = 'reorganized_BACH/malignant'
dest_b = 'reorganized_BACH/benign'

# Create the destination directory if it doesn't exist
os.makedirs('reorganized_BACH', exist_ok=True)

os.makedirs(dest_m, exist_ok=True)

os.makedirs(dest_b, exist_ok=True)
# Copy all files from the source directory to the destination directory
for file_name in os.listdir(src_b):
    source_file = os.path.join(src_b, file_name)
    destination_file = os.path.join(dest_b, file_name)
    if os.path.isfile(source_file):
        shutil.copy(source_file, destination_file)

print(f"All files have been copied from '{src_b}' to '{dest_b}'.")

for cur_src_m in all_src_m:
  for file_name in os.listdir(cur_src_m):
    source_file = os.path.join(cur_src_m, file_name)
    destination_file = os.path.join(dest_m, file_name)
    if os.path.isfile(source_file):
        shutil.copy(source_file, destination_file)

  print(f"All files have been copied from '{cur_src_m}' to '{dest_m}'.")

#transfer learning and final accuracy
import torch
from transformers import AutoModelForImageClassification, AutoFeatureExtractor
from torchvision.datasets import ImageFolder
from torchvision.transforms import Compose, Normalize, Resize, ToTensor
from torch.utils.data import DataLoader
from sklearn.metrics import confusion_matrix, precision_score, recall_score, f1_score, accuracy_score, roc_curve, matthews_corrcoef
from datasets import load_dataset

import numpy as np
import tensorflow as tf
from PIL import Image
from sklearn.model_selection import KFold


from transformers import AutoImageProcessor
from torchvision.transforms import RandomResizedCrop, Compose, Normalize, ToTensor

from tensorflow import keras
from tensorflow.keras import layers
from transformers import DefaultDataCollator


# Load the dataset using ImageFolder
dataset = load_dataset("imagefolder",data_dir='reorganized_BACH', split='train')
#dataset = ImageFolder(root='reorganized_idc', transform=transform)

#dataset = dataset.train_test_split(test_size=0.4)

dataset = dataset.train_test_split(test_size=0.2)


print(dataset["train"][0])

import numpy as np
import tensorflow as tf
from PIL import Image


def convert_to_tf_tensor(image: Image):
    np_image = np.array(image)
    tf_image = tf.convert_to_tensor(np_image)
    # `expand_dims()` is used to add a batch dimension since
    # the TF augmentation layers operates on batched inputs.
    return tf.expand_dims(tf_image, 0)


def preprocess_train(example_batch):
    """Apply train_transforms across a batch."""
    images = [
        train_data_augmentation(convert_to_tf_tensor(image.convert("RGB"))) for image in example_batch["image"]
    ]
    example_batch["pixel_values"] = [tf.transpose(tf.squeeze(image)) for image in images]
    return example_batch


def preprocess_val(example_batch):
    """Apply val_transforms across a batch."""
    images = [
        val_data_augmentation(convert_to_tf_tensor(image.convert("RGB"))) for image in example_batch["image"]
    ]
    example_batch["pixel_values"] = [tf.transpose(tf.squeeze(image)) for image in images]
    return example_batch

labels=['benign', 'malignant']
label2id, id2label= dict(), dict()

for i, label in enumerate(labels):
  label2id[label] = str(i)
  id2label[str(i)] = label


image_processor=AutoImageProcessor.from_pretrained(model_id)


normalize = Normalize(mean=image_processor.image_mean, std=image_processor.image_std)

size = (
    image_processor.size["shortest_edge"]
    if "shortest_edge" in image_processor.size
    else (image_processor.size["height"], image_processor.size["width"])
)

_transforms = Compose([RandomResizedCrop(size), ToTensor(), normalize])

def transforms(examples):
    examples["pixel_values"] = [_transforms(img.convert("RGB")) for img in examples["image"]]
    del examples["image"]
    return examples


size = (image_processor.size["height"], image_processor.size["width"])
dataset["train"].set_transform(preprocess_train)
dataset["test"].set_transform(preprocess_val)

dataset = dataset.with_transform(transforms)

data_collator = DefaultDataCollator()

#ranger class as implemented in Wright et al. 2021, "Ranger21: A synergistic deep learning optimizer"
import math
import torch
from torch.optim.optimizer import Optimizer

torch.cuda.empty_cache()

class Ranger(Optimizer):

	def __init__(self, params, lr=1e-3,  # lr
				 alpha=0.5, k=6, N_sma_threshhold=5,  # Ranger options
				 betas=(.95, 0.999), eps=1e-5, weight_decay=0,  # Adam options
				 use_gc=True, gc_conv_only=False
				 # Gradient centralization on or off, applied to conv layers only or conv + fc layers
				 ):

		# parameter checks
		if not 0.0 <= alpha <= 1.0:
			raise ValueError(f'Invalid slow update rate: {alpha}')
		if not 1 <= k:
			raise ValueError(f'Invalid lookahead steps: {k}')
		if not lr > 0:
			raise ValueError(f'Invalid Learning Rate: {lr}')
		if not eps > 0:
			raise ValueError(f'Invalid eps: {eps}')

		# parameter comments:
		# beta1 (momentum) of .95 seems to work better than .90...
		# N_sma_threshold of 5 seems better in testing than 4.
		# In both cases, worth testing on your dataset (.90 vs .95, 4 vs 5) to make sure which works best for you.

		# prep defaults and init torch.optim base
		defaults = dict(lr=lr, alpha=alpha, k=k, step_counter=0, betas=betas, N_sma_threshhold=N_sma_threshhold,
						eps=eps, weight_decay=weight_decay)
		super().__init__(params, defaults)

		# adjustable threshold
		self.N_sma_threshhold = N_sma_threshhold

		# look ahead params

		self.alpha = alpha
		self.k = k

		# radam buffer for state
		self.radam_buffer = [[None, None, None] for ind in range(10)]

		# gc on or off
		self.use_gc = use_gc

		# level of gradient centralization
		self.gc_gradient_threshold = 3 if gc_conv_only else 1

	def __setstate__(self, state):
		super(Ranger, self).__setstate__(state)

	def step(self, closure=None):
		loss = None

		# Evaluate averages and grad, update param tensors
		for group in self.param_groups:

			for p in group['params']:
				if p.grad is None:
					continue
				grad = p.grad.data.float()

				if grad.is_sparse:
					raise RuntimeError('Ranger optimizer does not support sparse gradients')

				p_data_fp32 = p.data.float()

				state = self.state[p]  # get state dict for this param

				if len(state) == 0:  # if first time to run...init dictionary with our desired entries
					# if self.first_run_check==0:
					# self.first_run_check=1
					# print("Initializing slow buffer...should not see this at load from saved model!")
					state['step'] = 0
					state['exp_avg'] = torch.zeros_like(p_data_fp32)
					state['exp_avg_sq'] = torch.zeros_like(p_data_fp32)

					# look ahead weight storage now in state dict
					state['slow_buffer'] = torch.empty_like(p.data)
					state['slow_buffer'].copy_(p.data)

				else:
					state['exp_avg'] = state['exp_avg'].type_as(p_data_fp32)
					state['exp_avg_sq'] = state['exp_avg_sq'].type_as(p_data_fp32)

				# begin computations
				exp_avg, exp_avg_sq = state['exp_avg'], state['exp_avg_sq']
				beta1, beta2 = group['betas']

				# GC operation for Conv layers and FC layers
				if grad.dim() > self.gc_gradient_threshold:
					grad.add_(-grad.mean(dim=tuple(range(1, grad.dim())), keepdim=True))

				state['step'] += 1

				# compute variance mov avg
				exp_avg_sq.mul_(beta2).addcmul_(1 - beta2, grad, grad)
				# compute mean moving avg
				exp_avg.mul_(beta1).add_(1 - beta1, grad)

				buffered = self.radam_buffer[int(state['step'] % 10)]

				if state['step'] == buffered[0]:
					N_sma, step_size = buffered[1], buffered[2]
				else:
					buffered[0] = state['step']
					beta2_t = beta2 ** state['step']
					N_sma_max = 2 / (1 - beta2) - 1
					N_sma = N_sma_max - 2 * state['step'] * beta2_t / (1 - beta2_t)
					buffered[1] = N_sma
					if N_sma > self.N_sma_threshhold:
						step_size = math.sqrt(
							(1 - beta2_t) * (N_sma - 4) / (N_sma_max - 4) * (N_sma - 2) / N_sma * N_sma_max / (
										N_sma_max - 2)) / (1 - beta1 ** state['step'])
					else:
						step_size = 1.0 / (1 - beta1 ** state['step'])
					buffered[2] = step_size

				if group['weight_decay'] != 0:
					p_data_fp32.add_(-group['weight_decay'] * group['lr'], p_data_fp32)

				# apply lr
				if N_sma > self.N_sma_threshhold:
					denom = exp_avg_sq.sqrt().add_(group['eps'])
					p_data_fp32.addcdiv_(-step_size * group['lr'], exp_avg, denom)
				else:
					p_data_fp32.add_(-step_size * group['lr'], exp_avg)

				p.data.copy_(p_data_fp32)

				# integrated look ahead...
				# we do it at the param level instead of group level
				if state['step'] % group['k'] == 0:
					slow_p = state['slow_buffer']  # get access to slow param tensor
					slow_p.add_(self.alpha, p.data - slow_p)  # (fast weights - slow weights) * alpha
					p.data.copy_(slow_p)  # copy interpolated weights to RAdam param tensor

		return loss

import evaluate

accuracy = evaluate.load("accuracy")

import numpy as np

def compute_metrics(eval_pred):
    predictions, labels = eval_pred
    predictions = np.argmax(predictions, axis=1)
    accuracy_metric = accuracy.compute(predictions=predictions, references=labels)
    return accuracy_metric

#train the BACH transfer learning model
from transformers import get_scheduler
import pandas as pd
from transformers import AutoModelForImageClassification, TrainingArguments, Trainer

model = AutoModelForImageClassification.from_pretrained(
    model_id,
    num_labels=len(labels),
    id2label=id2label,
    label2id=label2id,
)
optimizer = Ranger(model.parameters(), lr=3e-5, weight_decay=.02)
training_args = TrainingArguments(
    output_dir="break_his_default_ranger_rotations_5_fold_1_transfer_learning_bach_final",
    remove_unused_columns=False,
    eval_strategy="epoch",
    save_strategy="epoch",
    learning_rate=1e-5,
    per_device_train_batch_size=8,
    gradient_accumulation_steps=2,
    per_device_eval_batch_size=8,
    num_train_epochs=100,
    warmup_ratio=0.2,
    logging_steps=10,
    load_best_model_at_end=False,
    metric_for_best_model="accuracy",
    push_to_hub=True,
    hub_private_repo=True,
    hub_model_id="your huggingface model path and id"
)

training_steps = training_args.num_train_epochs * len(dataset["train"]) // training_args.per_device_train_batch_size
lr_scheduler = get_scheduler(
    name="cosine",
    optimizer=optimizer,
    num_warmup_steps=training_steps//6
    ,
    num_training_steps=training_steps
)

trainer = Trainer(
    model=model,
    args=training_args,
    data_collator=data_collator,
    train_dataset=dataset["train"],
    eval_dataset=dataset["test"],
    optimizers=(optimizer, lr_scheduler),
    tokenizer=image_processor,
    compute_metrics=compute_metrics,
)

trainer.train()

res_log = pd.DataFrame(trainer.state.log_history)

#load base model for either transfer learning or accuracy check
import torch
from transformers import AutoModelForImageClassification, AutoFeatureExtractor
from torchvision.datasets import ImageFolder
from torchvision.transforms import Compose, Normalize, Resize, ToTensor
from torch.utils.data import DataLoader
from sklearn.metrics import confusion_matrix, precision_score, recall_score, f1_score, accuracy_score, roc_curve, matthews_corrcoef

# Load the pre-trained model and feature extractor

model_id="sloshywings/breakhis-model-ranger-rotations-5-fold-1-final"

print(model_id)
model = AutoModelForImageClassification.from_pretrained(model_id)
feature_extractor = AutoFeatureExtractor.from_pretrained(model_id)

# Define transformations
transform = Compose([
    Resize((224, 224)),
    ToTensor(),
    Normalize(mean=feature_extractor.image_mean, std=feature_extractor.image_std)
])

#!mkdir /content/IDC_dataset/
#This portion of the code assumes that the IDC dataset zip file was unpacked as described in the following command at the current level of directory: unzip /content/drive/MyDrive/breast_histology/IDC_dataset.zip -d /content/IDC_dataset/

#Generate dataset for IDC
import os
import shutil

# Path to the root directory containing the numbered folders
root_dir = 'IDC_dataset/'

# Paths for the new directories to be created
benign_dir = os.path.join('reorganized_idc/benign')
malignant_dir = os.path.join('reorganized_idc/malignant')

os.makedirs('reogranized_idc/', exist_ok=True)

# Create the new directories if they don't exist
os.makedirs(benign_dir, exist_ok=True)
os.makedirs(malignant_dir, exist_ok=True)

# Iterate over each numbered folder in the root directory
for folder in os.listdir(root_dir):
    folder_path = os.path.join(root_dir, folder)

    # Check if it's a directory
    if os.path.isdir(folder_path):
        # Define the paths for the 0 (benign) and 1 (malignant) folders
        benign_folder_path = os.path.join(folder_path, '0')
        malignant_folder_path = os.path.join(folder_path, '1')

        # If the benign folder exists, copy its contents to the new benign directory
        if os.path.exists(benign_folder_path):
            for file_name in os.listdir(benign_folder_path):
                shutil.copy(os.path.join(benign_folder_path, file_name), benign_dir)

        # If the malignant folder exists, copy its contents to the new malignant directory
        if os.path.exists(malignant_folder_path):
            for file_name in os.listdir(malignant_folder_path):
                shutil.copy(os.path.join(malignant_folder_path, file_name), malignant_dir)

#transfer learning and final accuracy
import torch
from transformers import AutoModelForImageClassification, AutoFeatureExtractor
from torchvision.datasets import ImageFolder
from torchvision.transforms import Compose, Normalize, Resize, ToTensor
from torch.utils.data import DataLoader
from sklearn.metrics import confusion_matrix, precision_score, recall_score, f1_score, accuracy_score, roc_curve, matthews_corrcoef
from datasets import load_dataset

import numpy as np
import tensorflow as tf
from PIL import Image
from sklearn.model_selection import KFold


from transformers import AutoImageProcessor
from torchvision.transforms import RandomResizedCrop, Compose, Normalize, ToTensor

from tensorflow import keras
from tensorflow.keras import layers
from transformers import DefaultDataCollator


# Load the dataset using ImageFolder
dataset = load_dataset("imagefolder",data_dir='reorganized_idc', split='train')

dataset = dataset.train_test_split(test_size=0.4)


print(dataset["train"][0])

import numpy as np
import tensorflow as tf
from PIL import Image


def convert_to_tf_tensor(image: Image):
    np_image = np.array(image)
    tf_image = tf.convert_to_tensor(np_image)
    # `expand_dims()` is used to add a batch dimension since
    # the TF augmentation layers operates on batched inputs.
    return tf.expand_dims(tf_image, 0)


def preprocess_train(example_batch):
    """Apply train_transforms across a batch."""
    images = [
        train_data_augmentation(convert_to_tf_tensor(image.convert("RGB"))) for image in example_batch["image"]
    ]
    example_batch["pixel_values"] = [tf.transpose(tf.squeeze(image)) for image in images]
    return example_batch


def preprocess_val(example_batch):
    """Apply val_transforms across a batch."""
    images = [
        val_data_augmentation(convert_to_tf_tensor(image.convert("RGB"))) for image in example_batch["image"]
    ]
    example_batch["pixel_values"] = [tf.transpose(tf.squeeze(image)) for image in images]
    return example_batch

labels=['benign', 'malignant']
label2id, id2label= dict(), dict()

for i, label in enumerate(labels):
  label2id[label] = str(i)
  id2label[str(i)] = label


image_processor=AutoImageProcessor.from_pretrained(model_id)


normalize = Normalize(mean=image_processor.image_mean, std=image_processor.image_std)

size = (
    image_processor.size["shortest_edge"]
    if "shortest_edge" in image_processor.size
    else (image_processor.size["height"], image_processor.size["width"])
)

_transforms = Compose([RandomResizedCrop(size), ToTensor(), normalize])

def transforms(examples):
    examples["pixel_values"] = [_transforms(img.convert("RGB")) for img in examples["image"]]
    del examples["image"]
    return examples


size = (image_processor.size["height"], image_processor.size["width"])
dataset["train"].set_transform(preprocess_train)
dataset["test"].set_transform(preprocess_val)

dataset = dataset.with_transform(transforms)

data_collator = DefaultDataCollator()

#ranger class as implemented in Wright et al. 2021, "Ranger21: A synergistic deep learning optimizer"
import math
import torch
from torch.optim.optimizer import Optimizer

torch.cuda.empty_cache()

class Ranger(Optimizer):

	def __init__(self, params, lr=1e-3,  # lr
				 alpha=0.5, k=6, N_sma_threshhold=5,  # Ranger options
				 betas=(.95, 0.999), eps=1e-5, weight_decay=0,  # Adam options
				 use_gc=True, gc_conv_only=False
				 # Gradient centralization on or off, applied to conv layers only or conv + fc layers
				 ):

		# parameter checks
		if not 0.0 <= alpha <= 1.0:
			raise ValueError(f'Invalid slow update rate: {alpha}')
		if not 1 <= k:
			raise ValueError(f'Invalid lookahead steps: {k}')
		if not lr > 0:
			raise ValueError(f'Invalid Learning Rate: {lr}')
		if not eps > 0:
			raise ValueError(f'Invalid eps: {eps}')

		# parameter comments:
		# beta1 (momentum) of .95 seems to work better than .90...
		# N_sma_threshold of 5 seems better in testing than 4.
		# In both cases, worth testing on your dataset (.90 vs .95, 4 vs 5) to make sure which works best for you.

		# prep defaults and init torch.optim base
		defaults = dict(lr=lr, alpha=alpha, k=k, step_counter=0, betas=betas, N_sma_threshhold=N_sma_threshhold,
						eps=eps, weight_decay=weight_decay)
		super().__init__(params, defaults)

		# adjustable threshold
		self.N_sma_threshhold = N_sma_threshhold

		# look ahead params

		self.alpha = alpha
		self.k = k

		# radam buffer for state
		self.radam_buffer = [[None, None, None] for ind in range(10)]

		# gc on or off
		self.use_gc = use_gc

		# level of gradient centralization
		self.gc_gradient_threshold = 3 if gc_conv_only else 1

	def __setstate__(self, state):
		super(Ranger, self).__setstate__(state)

	def step(self, closure=None):
		loss = None

		# Evaluate averages and grad, update param tensors
		for group in self.param_groups:

			for p in group['params']:
				if p.grad is None:
					continue
				grad = p.grad.data.float()

				if grad.is_sparse:
					raise RuntimeError('Ranger optimizer does not support sparse gradients')

				p_data_fp32 = p.data.float()

				state = self.state[p]  # get state dict for this param

				if len(state) == 0:  # if first time to run...init dictionary with our desired entries
					# if self.first_run_check==0:
					# self.first_run_check=1
					# print("Initializing slow buffer...should not see this at load from saved model!")
					state['step'] = 0
					state['exp_avg'] = torch.zeros_like(p_data_fp32)
					state['exp_avg_sq'] = torch.zeros_like(p_data_fp32)

					# look ahead weight storage now in state dict
					state['slow_buffer'] = torch.empty_like(p.data)
					state['slow_buffer'].copy_(p.data)

				else:
					state['exp_avg'] = state['exp_avg'].type_as(p_data_fp32)
					state['exp_avg_sq'] = state['exp_avg_sq'].type_as(p_data_fp32)

				# begin computations
				exp_avg, exp_avg_sq = state['exp_avg'], state['exp_avg_sq']
				beta1, beta2 = group['betas']

				# GC operation for Conv layers and FC layers
				if grad.dim() > self.gc_gradient_threshold:
					grad.add_(-grad.mean(dim=tuple(range(1, grad.dim())), keepdim=True))

				state['step'] += 1

				# compute variance mov avg
				exp_avg_sq.mul_(beta2).addcmul_(1 - beta2, grad, grad)
				# compute mean moving avg
				exp_avg.mul_(beta1).add_(1 - beta1, grad)

				buffered = self.radam_buffer[int(state['step'] % 10)]

				if state['step'] == buffered[0]:
					N_sma, step_size = buffered[1], buffered[2]
				else:
					buffered[0] = state['step']
					beta2_t = beta2 ** state['step']
					N_sma_max = 2 / (1 - beta2) - 1
					N_sma = N_sma_max - 2 * state['step'] * beta2_t / (1 - beta2_t)
					buffered[1] = N_sma
					if N_sma > self.N_sma_threshhold:
						step_size = math.sqrt(
							(1 - beta2_t) * (N_sma - 4) / (N_sma_max - 4) * (N_sma - 2) / N_sma * N_sma_max / (
										N_sma_max - 2)) / (1 - beta1 ** state['step'])
					else:
						step_size = 1.0 / (1 - beta1 ** state['step'])
					buffered[2] = step_size

				if group['weight_decay'] != 0:
					p_data_fp32.add_(-group['weight_decay'] * group['lr'], p_data_fp32)

				# apply lr
				if N_sma > self.N_sma_threshhold:
					denom = exp_avg_sq.sqrt().add_(group['eps'])
					p_data_fp32.addcdiv_(-step_size * group['lr'], exp_avg, denom)
				else:
					p_data_fp32.add_(-step_size * group['lr'], exp_avg)

				p.data.copy_(p_data_fp32)

				# integrated look ahead...
				# we do it at the param level instead of group level
				if state['step'] % group['k'] == 0:
					slow_p = state['slow_buffer']  # get access to slow param tensor
					slow_p.add_(self.alpha, p.data - slow_p)  # (fast weights - slow weights) * alpha
					p.data.copy_(slow_p)  # copy interpolated weights to RAdam param tensor

		return loss

import evaluate

accuracy = evaluate.load("accuracy")

import numpy as np

def compute_metrics(eval_pred):
    predictions, labels = eval_pred
    predictions = np.argmax(predictions, axis=1)
    accuracy_metric = accuracy.compute(predictions=predictions, references=labels)
    return accuracy_metric

#train the BACH transfer learning model
from transformers import get_scheduler
import pandas as pd
from transformers import AutoModelForImageClassification, TrainingArguments, Trainer

model = AutoModelForImageClassification.from_pretrained(
    model_id,
    num_labels=len(labels),
    id2label=id2label,
    label2id=label2id,
)
optimizer = Ranger(model.parameters(), lr=3e-5, weight_decay=.02)
training_args = TrainingArguments(
    output_dir="break_his_default_ranger_rotations_5_fold_1_transfer_learning_idc_final",
    remove_unused_columns=False,
    eval_strategy="epoch",
    save_strategy="epoch",
    learning_rate=1e-5,
    per_device_train_batch_size=8,
    gradient_accumulation_steps=2,
    per_device_eval_batch_size=8,
    num_train_epochs=100,
    warmup_ratio=0.2,
    logging_steps=10,
    load_best_model_at_end=False,
    metric_for_best_model="accuracy",
    push_to_hub=True,
    hub_private_repo=True,
    hub_model_id="your huggingface model id and path"
)

training_steps = training_args.num_train_epochs * len(dataset["train"]) // training_args.per_device_train_batch_size
lr_scheduler = get_scheduler(
    name="cosine",
    optimizer=optimizer,
    num_warmup_steps=training_steps//6
    ,
    num_training_steps=training_steps
)

trainer = Trainer(
    model=model,
    args=training_args,
    data_collator=data_collator,
    train_dataset=dataset["train"],
    eval_dataset=dataset["test"],
    optimizers=(optimizer, lr_scheduler),
    tokenizer=image_processor,
    compute_metrics=compute_metrics,
)

trainer.train()

res_log = pd.DataFrame(trainer.state.log_history)
```
