## Supplementary material for "High-Performance Classification of Breast Cancer Histopathological Images Using Fine-Tuned Vision Transformers on the BreakHis Dataset": Code: README

These scripts are modified, auto generated python scripts based on the notebook files run on Google Colab using cloud resources. They are included for anyone that wants to repeat results on a local machine. There are a few places where the code must be modified to include appropriate directory paths and api keys or tokens specific to the user. See comments in code for more details. For the original notebook files, see the python_notebook directory files.
